# High throughput mechanobiology: Force modulation of ensemble biochemical and cell-based assays

**DOI:** 10.1101/2020.05.05.065912

**Authors:** Ália dos Santos, Natalia Fili, David S. Pearson, Yukti Hari-Gupta, Christopher P. Toseland

## Abstract

Mechanobiology is focused on how the physical forces and the mechanical properties of proteins, cells and tissues contribute to physiology and disease. While the response of proteins and cells to mechanical stimuli is critical for function, the tools to probe these activities are typically restricted to single molecule manipulations. Here, we have developed a novel microplate reader assay to encompass mechanical measurements with ensemble biochemical and cellular assays, using a microplate lid modified with magnets. This configuration enables multiple static magnetic tweezers to function simultaneously across the microplate, thereby greatly increasing throughput. The broad applicability and versatility of our approach has been demonstrated through *in vitro* force-induced enzymatic activity and conformation changes, along with force-induced receptor activation and their downstream signalling pathways in live cells. Overall, our methodology allows for the first-time ensemble biochemical and cell-based assays to be performed under force, in high throughput format. This novel approach would substantially add to the mechano-biological toolbox and increase the availability of mechanobiology measurements.

## INTRODUCTION

Mechanobiology focuses on understanding how physical forces correlate with protein, cell and tissue dynamics and organization through mechano-transduction (Jansen et al., 2015). It is emerging that mechano-transduction affects almost all cellular processes, from cell-cell and cell-extracellular matrix adhesions to cytoskeletal architecture and gene expression (Jansen et al., 2015, Uhler and Shivashankar, 2017). In this manner, physical forces provide a mechanism to propagate signals within and between cells (Mohammed et al., 2019).

Specialized single molecule force measurements (such as atomic force microscopy, optical traps and magnetic tweezers) have been advancing rapidly over the past two decades and have revealed how force is used in biological systems, from individual proteins to complexes and from individual organelles to cells (Elosegui-Artola et al., 2017, Seo et al., 2016, Yao et al., 2016, Lherbette et al., 2017). More recently, these techniques have been combined with high-resolution single molecule fluorescence approaches and cell imaging to visualise dynamic systems under the controlled application of force (Cordova et al., 2014, Madariaga-Marcos et al., 2018, Newton et al., 2019, Swoboda et al., 2014). These measurements have started to probe cellular systems in order to dissect mechano-transduction pathways.

However, single molecule manipulation experiments are challenging due to the need for specialized equipment and technical expertise, along with the time to acquire statistically relevant data. Moreover, technical challenges typically limit the experimental design to elegant but minimalized systems, and assays must be optimized for single molecule conditions. For example, only low concentrations can be used and the reactants must not interfere with the optical detection. Conversely, commonly used ensemble biochemical assays are performed in readily available microplate readers in order to increase their throughput in determining biochemical constants. However, while off-line mechanical measurements have been performed in microplates (Spencer et al., 2016), real-time plate reader assays do not allow mechanical manipulation during measurements and, therefore, are not suitable for performing traditional, established biochemical assays in the presence of force. Therefore, in order to understand the role of force in biological systems, it is critical to fuse these two approaches.

To this end, we have developed a novel mechano-biology assay that couples the relative ease of ensemble biochemical assays with magnetic tweezer-based manipulations, in order to quantitatively study biological processes in the absence and presence of force. This is achieved through the use of magnets incorporated into the lid of a microplate, allowing the assays to be performed in fluorescent microplate readers. Force manipulation of the sample occurs though the tension applied by the magnets to paramagnetic particles attached to the biomolecules of interest. The force applied to the sample can be easily tuned by changing the size of the magnets or adjusting the distance between magnet and sample.

## RESULTS AND DISCUSSION

### Assay Microplate Design

The assay is based around the use of neodymium magnets incorporated into a 3D printed microplate lid and biological samples coupled to paramagnetic particles, essentially mimicking multiple magnetic tweezers across the microplate (Figure 1A/B). Using a 24-well microplate as an example, a magnet configuration, consisting of two 5 mm cube magnets, as used in a magnetic tweezer assay, can be incorporated onto the lid and inserted into each of the microplate wells. The applied force is dependent upon (i) the distance between the magnets and the sample, which can be varied by the size of spacers within the lid (Figure 1A), (ii) the distance between the magnets and (iii) the size of the paramagnetic particles used(Yu et al., 2014). Therefore, by manipulating these properties, we can apply various levels of force across the microplate. Then, we can calculate an apparent force applied to the biological sample on the basis of the known properties’ values (Yu et al., 2014), as demonstrated in Figure 1C. The term apparent force is used to differentiate the absolute force measurements performed under single molecule manipulation conditions. This difference occurs due to the inclusion of multiple attachments to a bead within our ensemble data, which would lead to higher exerted forces, versus the selection of only single bead attachments. However, assuming bead attachment is stochastic, the same number and single and multiple attachments would occur in each microplate well. Therefore, this is a consistent error in our estimation of exerted force.

**Figure 1:**
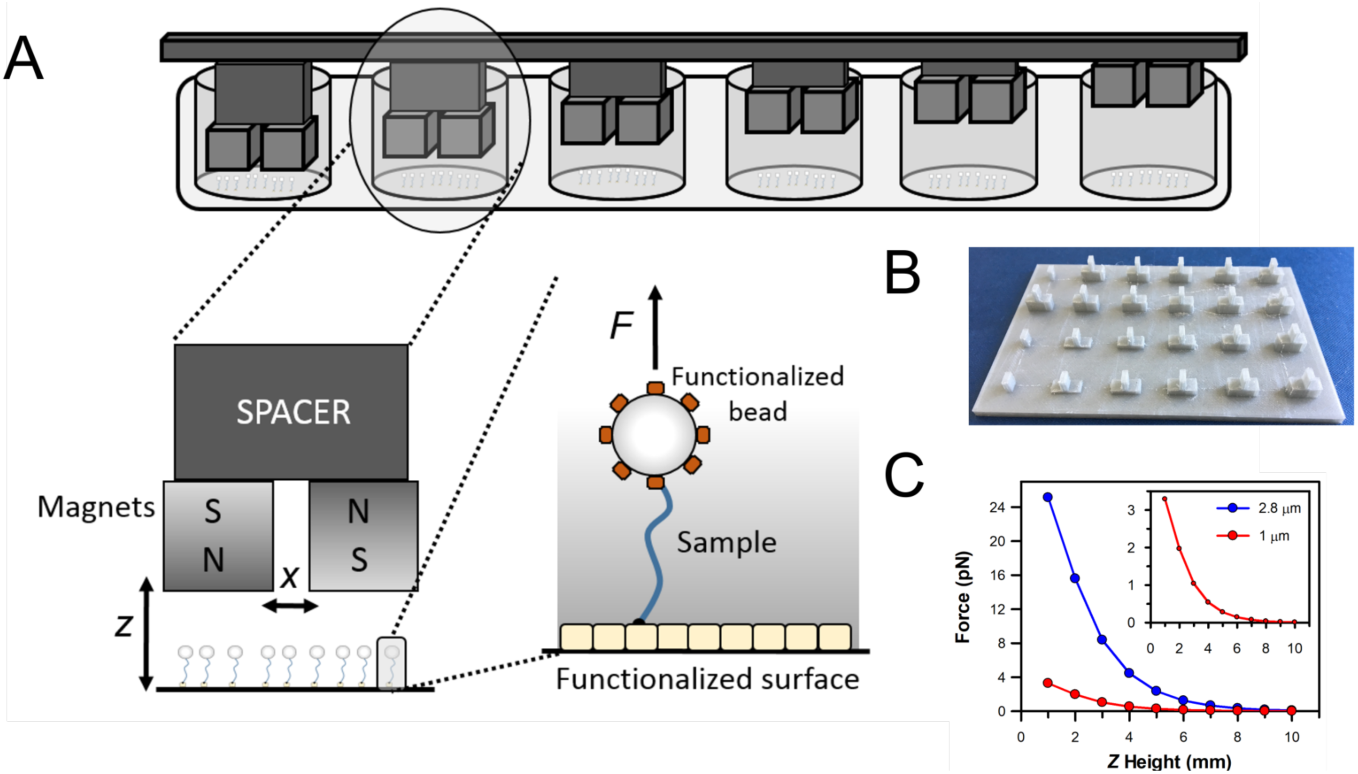
Multiplexed magnetic tweezers in a microplate. **(A)** Schematic of the assay’s format detailing the position and arrangement of neodymium cube magnets within a microplate-based assay. In this format, paramagnetic particles are coupled to the biological molecules of interest, which are in turn attached to the microplate surface. The position of the magnets above the surface (*Z* axis) is defined by spacers incorporated into the 3D printed lid **(B)**. The divider which sits on top of the spacers separates the magnets in the *X* axis by 2 mm. **(C)** The force exerted upon the biological sample is defined by the size of the paramagnetic particle, the distance of *Z* above the sample and the spacing between the magnets in *X*. The force-distance relationship is plotted for the 1 μm and 2.8 μm particles used in these studies. The inset shows the high distance, low force range.

The assay is configured to be compatible with a fluorescent microplate plate reader. It therefore allows simultaneous multi-colour assays over a broad spectrum of wavelengths, as well as the monitoring of intrinsic fluorophores. Here, the measurements are confined to single point at the centre of the well where, as with magnetic tweezers, the magnetic field is uniform. The diverse applications of this approach are demonstrated here by monitoring force-induced enzymatic activity, protein conformation changes and force-dependent signalling pathways. To assess the force dependence of these readouts, we have varied the strength of the applied magnetic field by controlling the distance between the magnets and the sample, as well as the size of the paramagnetic beads. Overall, we have been able to induce biological responses through the modulation of force, whilst simultaneously observing the response through fluorescent microplate-based assays.

### Proof of Principle: Force modulation of protein folding

As a proof of principle, we first used a recombinant Förster resonance energy transfer (FRET) based force reporter (Figure 2A). This was constructed by fusing two fluorescent proteins (eGFP donor and mRFP acceptor) to either end of the tension sensitive peptide repeat from flagelliform, whereby the FRET signal reports upon the force acting across the sensor by unfolding the peptide repeats, as described by Grashoff *et al* (Grashoff et al., 2010). The sensor also carried an N-terminal biotinylation tag to enable surface immobilisation, with its C-terminus coupled to Protein-A Dynabeads through an antibody against GFP. Fluorescence spectra of GFP and RFP were recorded under different experimental formats and the relative FRET signal was calculated from the GFP and RFP emission peaks. The 0 pN measurement was performed in the absence of magnets and provides a maximal FRET signal in this experimental format (Figure 2B). The absence of Dynabeads or antibody resulted in a higher starting FRET value, suggesting a partial unfolding of the construct, or a change in its fluorescence properties, when coupled to the antibody/Dynabeads. Indeed, the FRET signal is similar to that found with the reporter free in solution.

**Figure 2:**
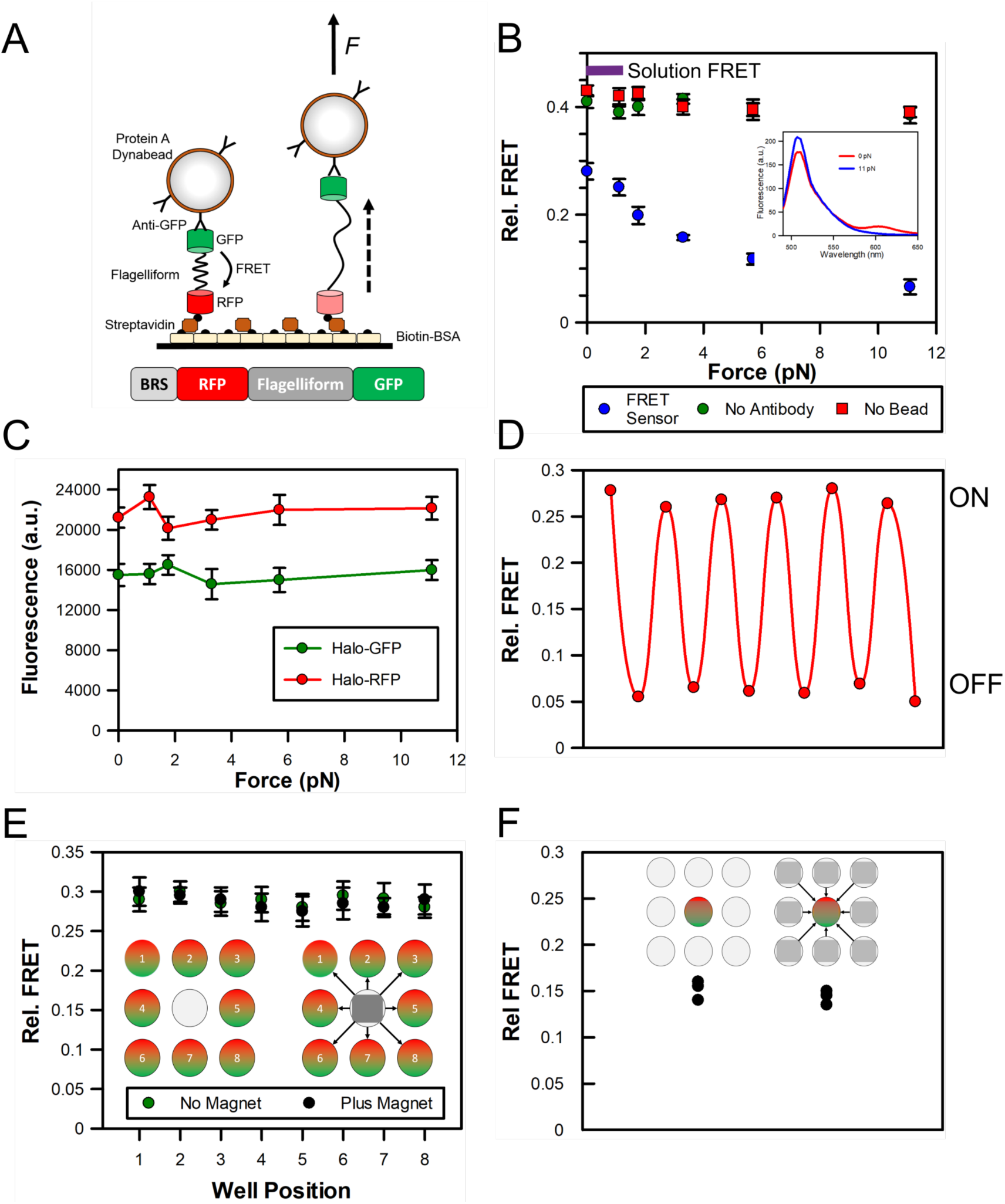
Proof-of-concept assay format monitoring protein unfolding. **(A)** Cartoon depicting the assay format monitoring protein unfolding through an eGFP-Flagelliform(8*GPGGA)-mRFP FRET sensor. The sensor was immobilised on the surface through an N-terminal biotinylation tag and was coupled to Protein-A Dynabeads through an antibody against GFP **(B)** Plot of Relative FRET signal obtained at different forces. The inset gives an example fluorescence spectra at 0 pN and 11 pN. Error bars represent SEM from 3 independent experiments. The purple bar indicates the relative FRET value of the sensor, free in solution. **(C)** Fluorescence intensity of eGFP-Flagelliform(8*GPGGA)-HaloTag (510 nm) and mRFP-Flagelliform(8*GPGGA)-HaloTag (605 nm) obtained at different forces. Error bars represent SEM from 3 independent experiments. **(D)** Example data showing alternations between high and low FRET states in the presence (ON) and absence (OFF) of 11 pN of force within a single microplate well. The oscillation is consistent with folding and unfolding of the flagelliform repeat. **(E-F)** Assess the impact of the magnetic fields upon surrounding microplate wells. **(E)** Cartoons depict the experimental setup. The FRET sensor was placed in 8 wells surrounding an empty central well. The Relative FRET signal was then determined for each well. The 11 pN magnet was positioned at the central well and the measurement was repeated. Error bars represent SEM from 3 independent experiments. **(F)** Cartoon depict the experimental setup. The FRET sensor was placed in a central well, exposed to a 6 pN magnet and the FRET signal was determined. 11 pN magnets were then placed in the surrounding 8 wells and the measurement was repeated. The experiment was performed in triplicate.

In the presence of the magnetic lid, (Figure 2B) there was an incremental decrease in FRET dependent upon the force applied (Figure 2B), as expected due to unfolding of the flagelliform peptide and the resulting increase in donor-acceptor distance. The absence of Dynabeads or antibody resulted in no changes in FRET signal in the presence of the magnets, indicating no further unfolding of the peptide. In order to ensure that the observed FRET signal was not due to intensity changed induced to the fluorescent proteins, two additional constructs were engineered: a BRS-mRFP-Flagelliform-Halo and a BRS-GFP-Flagelliform-Halo, where anti-Halo antibody was used to couple the constructs to the Dynabeads. Here, the magnet-induced protein unfolding did not cause any intensity changes within the individual fluorescent proteins (Figure 2C). Therefore, we are confident that the FRET changes arise solely from protein unfolding. This was further supported by the repeated cycles of folding and unfolding which were observed by removal and addition of the magnetic lid, respectively (Figure 2D).

The use of a microplate offers the advantage of high experimental throughput, since a single microplate would accommodate multiple magnetic tweezers conditions, compared to a single condition per coverslip in microscopy formats. However, this advantage could be compromised by the close proximity of the magnets in adjacent wells. To assess if there is cross-talk between the magnetic fields of neighbouring magnets within different microplate wells, we placed a single magnet pair in the centre of the microplate. We then used the FRET reporter to assess the effect of its magnetic field in the surrounding 8 wells which were devoid of magnets. As demonstrated in Figure 2E, in the absence of magnets in the central well, the FRET signal in the surrounding 8 wells was similar and consistent with the folded conformation of the peptide. Importantly, there is no change in signal when the magnets were placed in the central well, indicating that there is no cross-talk between adjacent microplate wells. In addition, we assessed whether surrounding magnets impact the magnetic field within an individual well. Here, the FRET reporter was placed within a central well and exposed to a single magnet to exert a moderate level of force (~6 pN) for partial unfolding (Figure 2F). The assay was then repeated in the presence of magnets in adjacent wells. Similarly, no difference in the FRET signals was observed, which supports the conclusion that there is no cross-talk between the wells.

This methodology is implemented in four types of biological assays to demonstrate the versatility of this novel tool: (a) force-induced conformation changes in a molecular motor, (b) the unwinding of DNA catalysed by a helicase, (c) the force-dependent binding of the single-stranded binding protein (SSB) to ssDNA and (d) a force-activated cell signalling pathway in live cells.

### Force-induced conformation changes and ligand binding

As a first application of our approach, we tested its potential in monitoring the force-induced conformation changes within a protein, using a similar assay format as above. The actin-based molecular motors, myosins, are regulated by numerous factors including intramolecular back-folding (Fili and Toseland, 2019). Myosin VI has been shown to exist in an auto-inhibited state, whereby the association of binding partners, such as NDP52, triggers unfolding and activation of the motor protein (Fili et al., 2020, Fili et al., 2017). As a molecular motor, myosin VI is a force-sensing (Hari-Gupta et al., 2020) and generating protein, therefore we investigated if forces can also unfold the protein which may act as a form of regulation.

To test this, we used a GFP-Myosin VI Tail-RFP construct, previously used to investigate intramolecular back-folding (Fili et al., 2017). The conformation sensor was immobilised to the microplate surface through N-terminal biotinylation and was coupled to the Dynabeads using an anti-RFP antibody (Figure 3A).

**Figure 3:**
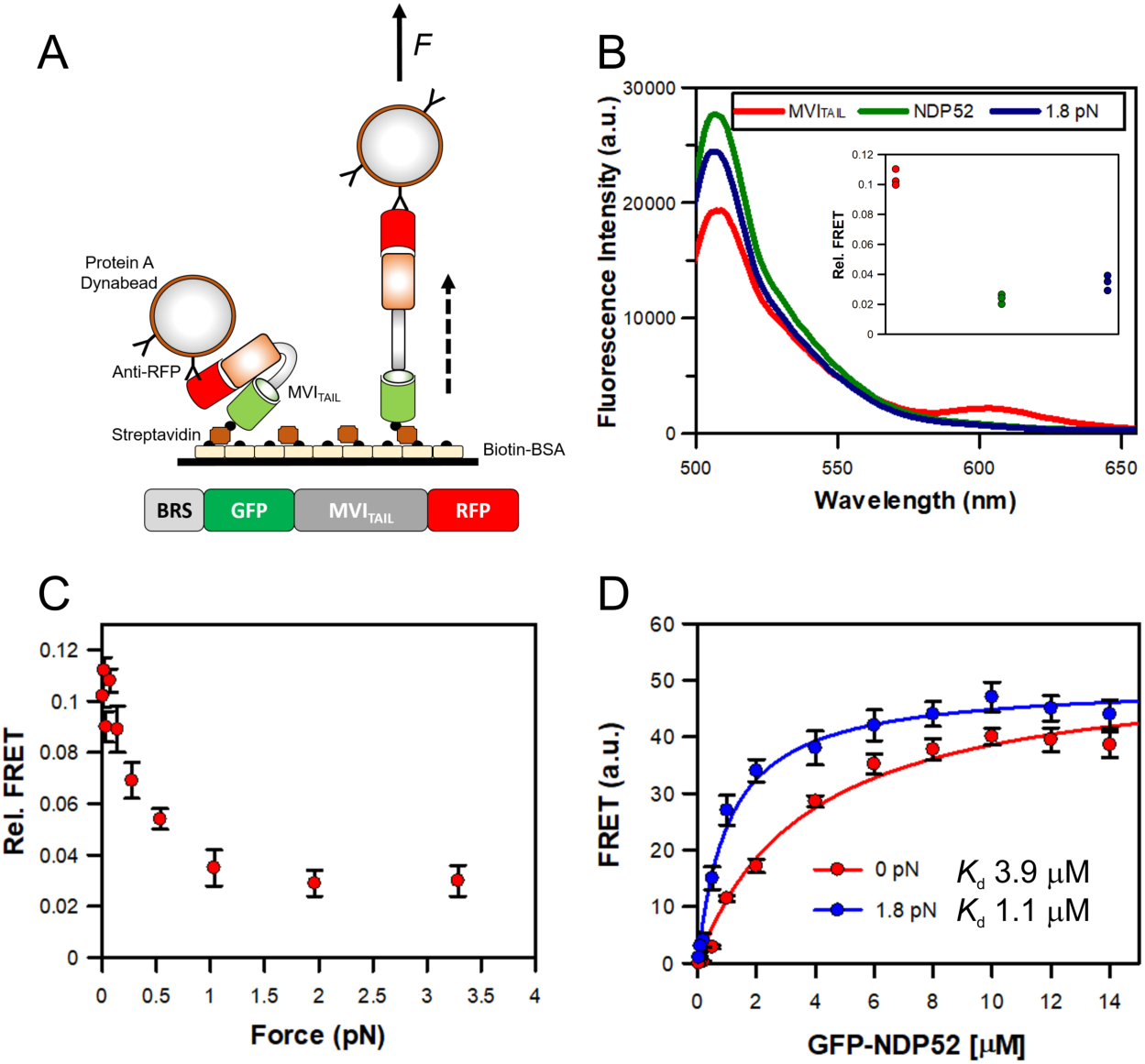
Force modulates myosin VI back-folding. **(A)** Cartoon depicting the assay format. The eGFP-MVI_TAIL_-mRFP construct was immobilised on the surface through an N-terminal biotinylation tag and it was coupled to Protein-A Dynabeads through an antibody against RFP. **(B)** Representative spectra showing GFP and RFP fluorescence in the absence of force, upon addition of 10 μM NDP52 or 1.8 pN force. **(C)** Plot of Relative FRET signal obtained at different forces. Error bars represent SEM from 3 independent experiments. **(D)** FRET Titration of GFP-NDP52 against BRS-MVI_TAIL_-mRFP under 0 pN (red) and 1.8 pN (blue). The myosin tail was immobilised using biotin and coupled to Protein-A Dynabeads through an antibody against RFP. Fitting the binding curve allowed *K*_d_ of 3.9 μM and 1.1 μM for 0 pN and 1.8 pN, respectively.

As shown previously, a FRET population corresponding to the back-folded myosin VI tail was observed and was lost following the addition of 10 µM NDP52, as unfolding occurred (Fili et al., 2017) (Figure 3B). Interestingly, the presence of 1.8 pN force also triggered unfolding of the tail domain. When the experiment was repeated for a range of forces, and the FRET signal did not change for forces above 1 pN (Figure 3C). Whilst outside the scope of this study, force within this range can be readily exerted upon other cytoskeletal motors through the application of load to the protein. It is therefore likely that force, generated by or acted upon the myosin enables unfolding while binding partners can stabilise this conformation.

The overriding benefit of this methodology is the ability to readily measure binding constants modulated by forces. Using the biological system above, we can measure the interaction between NDP52 and myosin VI in the presence and absence of force. Such an experimental approach will allow us to determine if NDP52 can bind to different conformations of myosin VI which will enable us to dissect the molecular mechanisms governing the myosin regulation. We modified the assay format to enable FRET to report upon the interaction between NDP52 and the myosin VI tail. Here, we used an RFP tag on the Myosin tail and GPF on NDP52. The tail was immobilised through biotin and coupled to dynabeads by anti-RFP. Titration of GFP-NDP52 in the absence of force yields an equilibrium dissociation constant, *K*_d_ 3.9 μM. This is essentially identical to the affinity calculated using traditional methods. The titration was then repeated in the presence of 1.8 pN force, which is enough to unfold the tail (Figure 3B). Binding was once again observed but a stronger interaction (*K*_d_ 1.1 μM) was measured. We can therefore conclude that NDP52 can interact with myosin VI in two conformations but that the unfolded state represents a more stable complex. It is therefore likely that NDP52 will bind to the folded complex and then trigger unfolding of the tail. Alternatively, force exerted by molecular cargo may have already unfolded the myosin and this complex would be stabilised by NDP52 binding.

These experiments represent the strength of this methodology whereby the effect of force on binding constants can be readily assessed.

### Force modulation of DNA Helicase activity

Another application of our approach is to assess the effect of force on the enzymatic activity of a protein of interest. To exemplify this, we used the bacterial DNA helicase PcrA. PcrA is a well-characterised ATP-dependent molecular motor, ideally suited for testing the feasibility of our experimental approach (Toseland et al., 2009). Furthermore, the motor is active when attached to glass surfaces and it can unwind linear DNA fragments (Chisty et al., 2013). As performed previously, we monitored DNA unwinding using a fluorescent single stranded DNA (ssDNA) biosensor (Fili et al., 2010, Fili et al., 2011). Biotinylated PcrA helicase was immobilised on optical glass bottom microplates through biotinylated BSA-streptavidin functionalisation (Figure 4A). A DNA fragment (1500 bp), with a single biotin tag at the 5’-end, was incubated with RepD to provide a loading site for PcrA helicase. The DNA fragment was then incubated with streptavidin Dynabeads, before blocking the remaining streptavidin binding sites on the bead with biotin-BSA. We used the same functionalisation process for both the surface and the beads, because they were performed independently.

**Figure 4:**
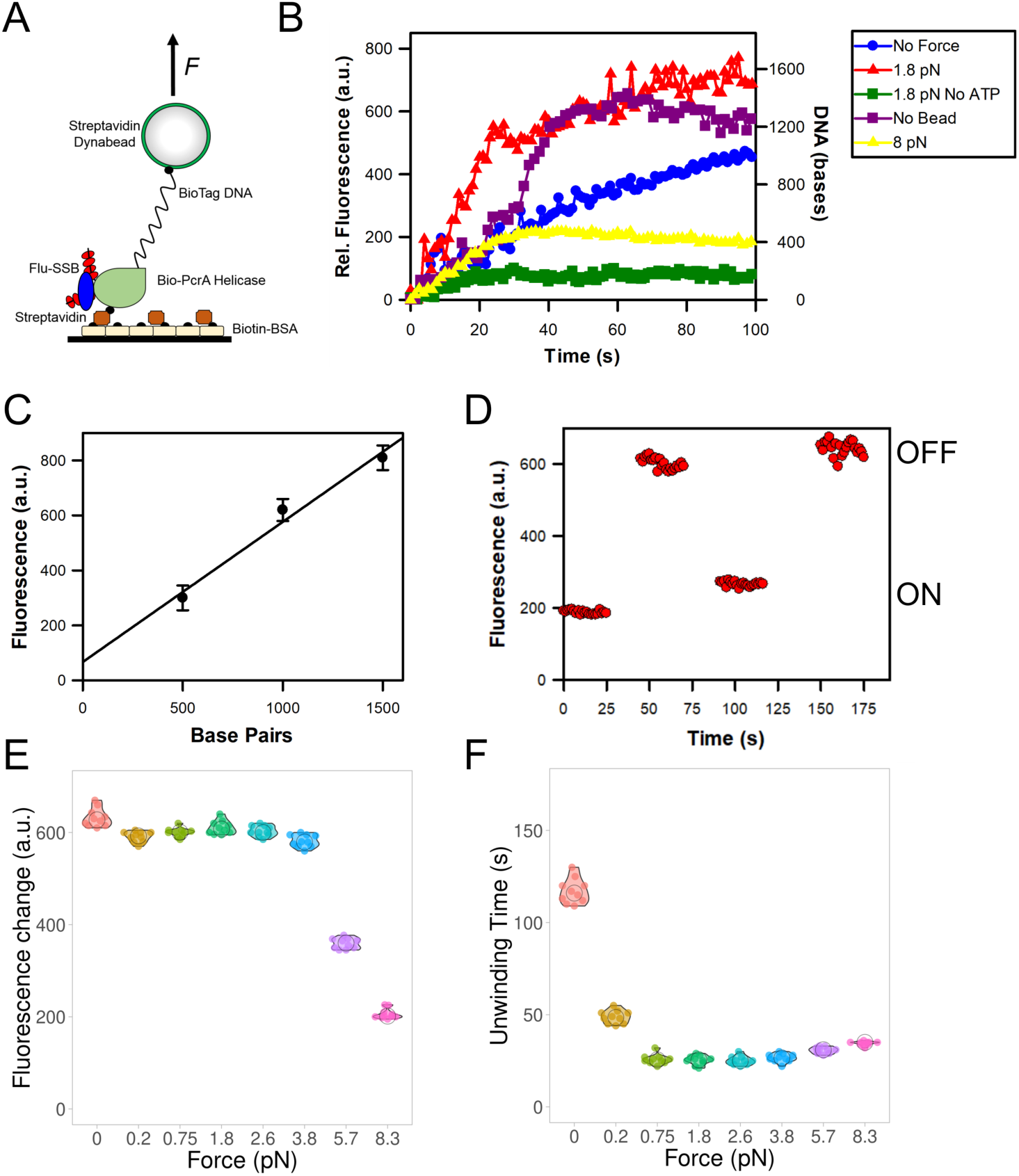
PcrA helicase activity on DNA held under tension. **(A)** Cartoon depicting the helicase unwinding assay. The helicase was immobilized on the surface, whilst RepD-loaded DNA was bound to the Dynabead, both through biotin-streptavidin interactions. Fluorescent SSB bound to the ssDNA product to report upon unwinding. **(B)** Representative traces of helicase activity. Helicase unwinding activity was monitored in real-time by recording the fluorescence change of MDCC-SSB binding to ssDNA. DNA unwinding in the presence of 8 pN force resulted in a lower fluorescence signal than the other unwinding experiments. Experiments were performed in the absence of Dynabead, magnets (No Force) and ATP. Fluorescence was converted to base pairs unwound using the calibration **(C)**, as described in the Methods. **(D)** The fluorescence signal from the DNA unwinding experiment performed at 8 pN could cycle through high (unwound) and low levels depending on the presence or absence of the magnet 8 pN magnet. **(E)** A summary of fluorescence change data for helicase unwinding experiments performed at the stated force levels. **(F)** A summary of unwinding time data for helicase unwinding experiments performed at the stated force levels.

As expected, the immobilised PcrA bound the DNA and then initiated unwinding in the presence of ATP. This was monitored in real-time using the fluorescent ssDNA biosensor, MDCC-SSB, whereby the biosensor bound the ssDNA product causing a fluorescence increase, following ATP-dependent DNA unwinding(Toseland and Webb, 2010). This assay allowed the helicase activity to be followed in the presence and absence of force (Figure 4B).

The fluorescence signal was calibrated against three lengths of DNA (Figure 4C) so that it is possible to measure unwinding in base-pairs per second. PcrA was able to unwind the DNA not bound to beads, as observed before (Chisty et al., 2013) with a lag-phase followed by an unwinding phase. PcrA unwound DNA at a rate of 37.5 bp s^−1^, while unwinding of DNA bound to the Dynabeads reduced unwinding to 11 bp s^−1^. The subsequent addition of 1.8 pN of force led to a six-fold increase in unwinding rate to 62 bp s^−1^. This is also two-fold higher than naked DNA and the trace does not contain the lag period before unwinding began. This suggests unwinding may begin more efficiently. Interestingly, the addition of 8 pN of force led to partial unwinding or a perturbed fluorescence signal. Assuming only partial unwinding occurred, then PcrA functioned at 16 bp s^−1^.

The addition of Dynabeads impeded DNA unwinding potentially through the addition of greater load on the DNA and/or increased surface interactions. The addition of the magnetic field would elongate the DNA, which would release surface interactions, uplifting the impediment. The overall increase in unwinding rate might reflect an rise in unwinding efficiency due to the elongated, linear DNA geometry. In this manner, the helicase would more readily separate the DNA strands. However, this did not occur when a higher force was applied; instead, in this case unwinding, or at least the measurement of unwinding was impeded. This may occur due to the displacement of the DNA from the helicase, to a force-induced stalling of the helicase or to the loss of the biosensor-DNA interactions. Interestingly, after removal of the high force magnet, a fluorescent signal consistent with complete unwinding was observed (Figure 4D). This was subsequently lost when the magnet was re-added. We therefore concluded that DNA unwinding did occur, however the biosensor-DNA interactions were perturbed by the high force, as investigated further below.

To demonstrate the throughput of this assay approach, we performed 10 helicase unwinding assays at 8 different force levels, where a measurement at each force level was performed simultaneously. We then calculate the maximum fluorescence change (Figure 4E) and the corresponding unwinding time, taken as the time to reach the maximum fluorescent signal (Figure 4F). Where the unwinding extent, rather than time, is important, it is possible to run the experiment offline and then scan the entire microplate. This level of throughput cannot be achieved with the corresponding tweezer assays. We find that each experiment was highly reproducible to approximately 10 %. Consistent with the data above, forces above 4 pN led to a decreased fluorescent signal. Moreover, unwinding time decreased as force was applied. These data highlight the potential of our approach to assess the effect of force on enzymatic activities. While magnetic tweezer experiments can measure unwinding events in explicit details, this approach allows to simultaneously record multiple DNA lengths under various conditions (ATP concentrations, salts, other protein factors). Conversely, single molecules measurements would require multiple coverslips per condition, thereby slowing the throughput and depth of the investigation.

### Protein-DNA interactions show a force dependence

Having monitored DNA unwinding by PcrA with the biosensor MDCC-SSB, we found that under high-force conditions there was a loss of fluorescent signal. As mentioned above, our data suggested that the interactions between SSB and DNA is force-dependent. To assess the nature of the SSB-ssDNA interaction, DNA substrates were generated with a 5’-digoxigenin and 3’-biotin tags to enable coupling of the substrate to the beads and surface, respectively (Figure 5A).

**Figure 5:**
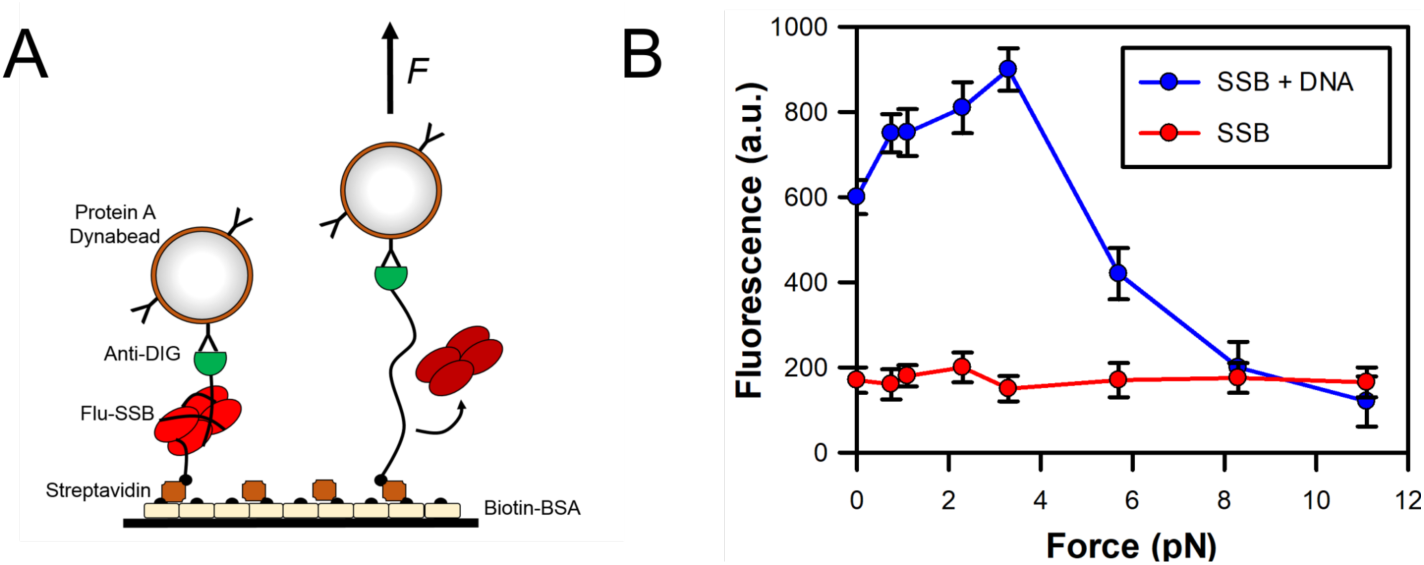
Force dependent interactions of SSB protein with ssDNA. **(A)** Cartoon showing the SSB interaction with ssDNA. ssDNA produced by PcrA unwinding was bound to the surface and paramagnetic bead using biotin and digoxigenin coupling, respectively. Fluorescent SSB was then added to the well and allowed to interact with the DNA. **(B)** The fluorescence from MDCC-SSB was recorded (at 475 nm) at various forces. Measurements were also recorded in the absence of DNA to provide the background signal generated by free SSB. SSB binding to ssDNA leads to a three-fold increase in fluorescence. Error bars represent SEM from 3 independent experiments.

The PcrA helicase was used to generate the ssDNA before the magnets were applied.

We have been able to observe that forces up to 4 pN led to an increase in fluorescence signal suggesting there was enhanced SSB binding onto ssDNA (Figure 5B). However, increasing the force above 4 pN led to a decrease in the fluorescence signal, with background levels being reached at 8 pN. This was consistent with the striping of SSB from the ssDNA, as indicated by Figure 4B and 4D and as proposed by Zhou *et al* (Zhou et al., 2011).

### Force modulation of cell signalling

Finally, in order to demonstrate the applicability of our tool to study mechano-transduction in live cells, we used a U2OS stable cell line expressing a SNAP-tagged human Notch1-Gal4 and a UAS-GAL4 mCherry reporter, which was previously developed for investigating the mechanical response of the Notch receptor (Seo et al., 2016). In that study, force applied by a magnetic tweezer microscope to the Notch receptor activated downstream signalling, which led to the expression and nuclear localisation of the mCherry reporter (Figure 6A). We set out to reproduce these experiments in a microplate reader format. U2OS cells were seeded onto a sterile microplate and allowed to grow to 50% confluency. The extracellular region of SNAP-tagged Notch1 receptors were first labelled with biotinylated SNAP ligand and then allowed to attach to streptavidin-coated paramagnetic beads for 1 hr, before the magnetic lid was applied. The reporter expression was then monitored by fluorescence for 24 hrs (Figure 6B). Applied force of 0.8 pN led to an increase in reporter expression by 14 hrs. This expression was 4-fold higher than that observed in the absence of magnet or Dynabead, thereby demonstrating mechanical activation of the Notch receptor pathway.

**Figure 6:**
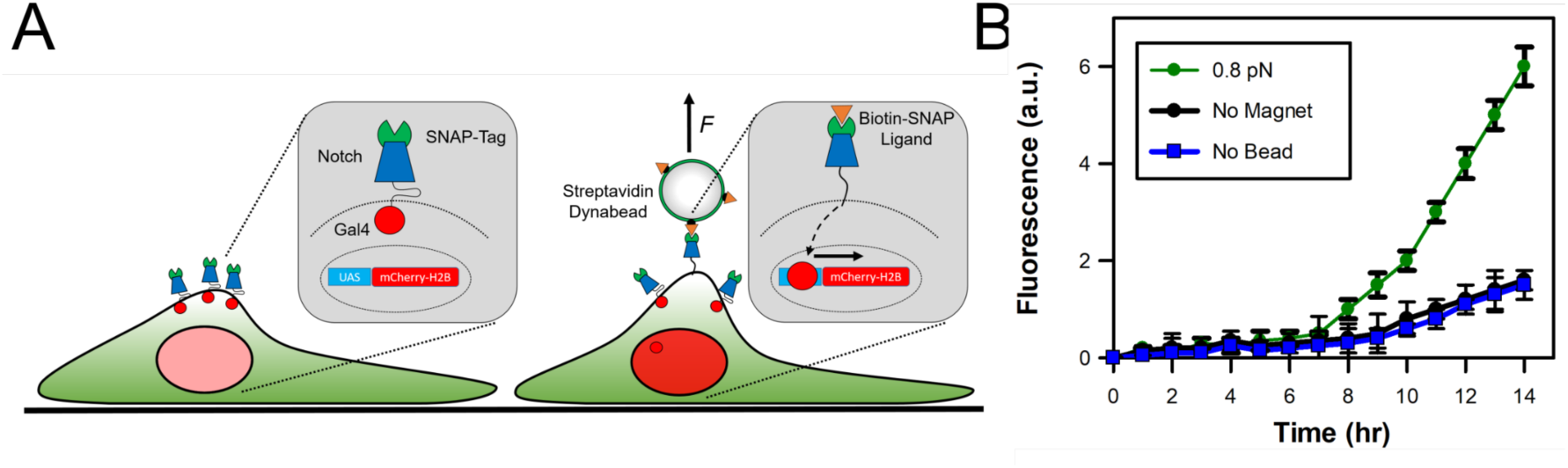
Force-activation Notch signalling. **(A)** Cartoon detailing the set-up of the live cell assay in order to monitor the force-induced activation of the notch receptor. The modified receptor was coupled to streptavidin-coated paramagnetic beads through a SNAP-biotin ligand. Force across the receptor led to cleavage of the C-terminal domain. In this instance, the C-terminal domain was replaced with Gal4 to act as a transcriptional reporter by driving the expression of mCherry-H2B. **(B)** The fluorescence emitting from the reporter was monitored in real-time. The fluorescence signal was dependent upon the presence of force and paramagnetic particles. Error bars represent SEM from 3 independent experiments.

This approach provides a controlled mechanism to activate a receptor in a high-throughput format therefore it would be possible to initiate drug screens against such pathway. No other methodology provides this capability and throughput.

### Conclusions

In summary, we have developed an easy-to-use, widely applicable tool for ensemble force-based measurements in a microplate format using fluorescent microplate readers. This methodology facilitates mechanobiology measurements in laboratories without the requirement for dedicated single molecule imaging facilities. However, we do not have the force accuracy found within single molecule measurements and therefore this approach does not aim to compete with these methodologies.

We have exemplified our technology through four different methods representing protein conformation changes, enzymatic activity, protein-ligand interactions and live cell reporter assays to demonstrate the versatility of the tool. Overall, this tool can induce biological responses through the modulation of force within novel assay formats. With this technology, it is now possible to quantitatively study a broad range of biological processes using established ensemble assays in a mechanical context, both *in vitro* and in cells, or tissues. The approach enables multiple conditions to be tested simultaneously, allowing high throughput force-induced measurements. Moreover, the microplate design provides high flexibility in the experimental format, ranging from 6-well to 96-well plate. Therefore, our approach is compatible with biochemical and cell-based assays through to advanced high-throughput drug screens. Furthermore, the microplate design is compatible with all microscopy-based formats, thereby enabling all imaging platforms to be combined with mechanical measurements and therefore expanding its range of applicability. This novel approach would substantially add to the mechano-biological toolbox available to study the impact of force in biological processes.

## METHODS

### Materials

Unless stated otherwise, all reagents were purchased from Sigma Aldrich (UK). All oligonucleotides were purchased from Integrated DNA Technologies (Belgium). Dynabeads were purchased from Thermo Fisher Scientific. MDCC-SSB was prepared as described in (Cook et al., 2018), BioPcrA was prepared as described in (Chisty et al., 2013) and RepD was prepared as described in (Toseland et al., 2009). BRS-eGFP-Myosin VI Tail-mRFP, NDP52 and GFP-NDP52 was prepared as described in (Fili et al., 2017).

Different lengths of linear DNA substrates were generated by PCR using the Hot start Phusion polymerase (New England Biolabs), as described previously based on the pCER*ori*D (Chisty et al., 2013). Primers containing 5’ or 3’ modifications for Biotin-TEG or digoxigenin were used to construct the substrates.

BRS-mRFP-Flagelliform(8*GPGGA)-eGFP, BRS-mRFP-Flagelliform(8*GPGGA)-HaloTag, BRS-eGFP-Flagelliform(8*GPGGA)-HaloTag, BRS-Myosin VI Tail-mRFP synthetic constructs were purchased from GeneArt (Thermo Fisher Scientific). Recombinant constructs were co-expressed with BirA in *E.coli* BL21 DE3 cells (Invitrogen) in Luria Bertani media supplemented with 50 mM biotin. Proteins were purified by affinity chromatography (HisTrap FF, GE Healthcare). The purest fractions were further purified through a Superdex 200 16/600 column (GE Healthcare).

The UAS-Gal4 reporter U2OS cells, which stably expressed SNAP-hN1-Gal4 and the H2B-mCherry fluorescence reporter sequence, were kindly gifted by Young-Wook Jun at UCSF.

### 3D printed lid Magnetic Lid

The microplate lid was designed using SketchUp (https://www.sketchup.com) and was produced using Co-Polyester on an Ultimaker 3 printer. 5mm cube Neodymium N42 magnets (supermagnete) were attached to the pedestals.

### Calculation of force exerted by the magnetic lid

The force exerted upon a sample is dependent upon the size of particle, gap between magnetic poles and distance between magnets and particle. The force is calculated using the equations below, based on the analysis of (Yu et al., 2014).

For a 2.8 μm particle with a 2 mm gap between magnetic poles:

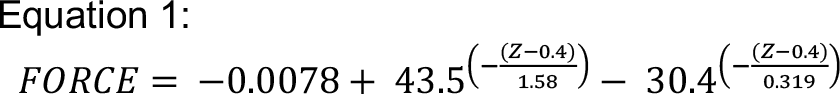

For a 1 μm particle with a 2 mm gap between magnetic poles:

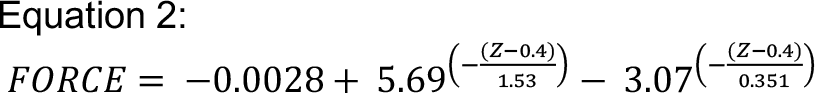

Where *Z* refers to the distance between the magnets and particle in millimeters.

### Surface modification for in vitro experiments

Glass-bottom microplates (Corning) were used in all assays. Surfaces were treated with 0.2mg/ml biotin-BSA in 50 mM Tris-HCl (pH7.5) and 50 mM NaCl for 15 min. Surfaces were then washed three times in buffer before adding 20 µg/ml streptavidin and incubating for 15 min. Surfaces were then washed three times with buffer.

### Fluorescence spectroscopy

All assays were performed on a BMG Labtech ClarioStar plate reader using bottom-optic configuration to facilitate the use of the magnetic lid.

### Experimental assays

#### FRET force sensor

Using the prepared surfaces, 100 nM biotin-mRFP-Flagelliform(8*GPGGA)-eGFP, biotin-mRFP-Flagelliform(8*GPGGA)-HaloTag, biotin-eGFP-Flagelliform(8*GPGGA)-HaloTag was added to the surface and allowed to incubate for 15 min at room temperature in 50 mM Tris-HCl (pH 7.5) and 50 mM NaCl. The surface was then washed three times in buffer. 1 mg Protein A dynabeads (2.8 μm) were prepared and loaded with 10 μg Anti-GFP antibody (Abcam ab290) or Anti-HaloTag (Promega G9281) according to the manufacturer’s instructions. 10 μg Dynabead-Antibody fusion was added to the surface and allowed to incubate for 30 min at room temperature. Surfaces were then washed three times in buffer. Fluorescence spectra corresponding to GFP and RFP were then recorded at 25 °C in the absence and presence of magnetic lid.

Data was processed to calculate Relative FRET.

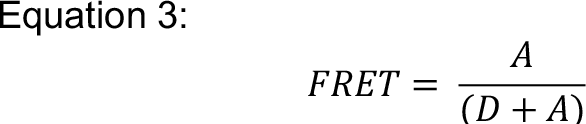

Where A is acceptor intensity (RFP 610 nm) and D is donor intensity (GFP 510 nm).

#### Force-induced conformation changes in Myosin VI

Using the prepared surfaces, 100 nM biotin-eGFP-Myosin VI Tail-mRFP or biotin-Myosin VI Tail-mRFP was added to the surface and allowed to incubate for 15 min at room temperature in 50 mM Tris-HCl (pH 7.5) and 50 mM NaCl. The surface was then washed three times in buffer. 1 mg Protein A dynabeads (2.8 μm) were prepared and loaded with 10 μg Anti-RFP antibody (Abcam ab62341) according to the manufacturer’s instructions. 10 μg Dynabead-Antibody fusion was added to the surface and allowed to incubate for 30 min at room temperature. Surfaces were then washed three times in buffer. Fluorescence spectra corresponding to GFP and RFP were then recorded at 25 °C in the absence and presence of magnetic lid, or in the presence of 10 μM NDP52. Data was processed to calculate Relative FRET using equation 3. For titrations, the fluorescence at 605 nm was measured during titration of GFP-NDP52. Background fluorescence was subtracted from the 605 nm intensity. The titration curves were fitted using equation 4:

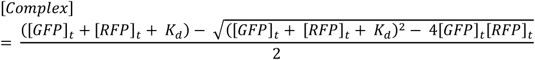

#### Surface-immobilized PcrA helicase assays with linear DNA

The streptavidin-coated surface was first treated with 10 nM bioPcrA in 50 mM Tris-HCl (pH 7.5), 150 mM NaCl, 3 mM MgCl_2_ and 1 mM DTT for 30 min. 1 mg MyOne Streptavidin C1 dynabeads (1 μm) were prepared according to the manufacturer’s instructions. 1 μg 1500 bp DNA substrate was incubated for 15 min at room temperature. Beads were washed three times in buffer. 30 μg dynabeads were added to each well. 500 nM RepD was then added to the well and incubated for 10 min. The unwinding was initiated with 1 mM ATP, supplemented with 100 nM MDCC-SSB. The reactions were performed at 25 C and MDCC fluorescence (ex. 430nm and em. 470 nm) was monitored in real-time in the presence and absence of the magnetic lid.

To calibrate the fluorescence signal, three different lengths of DNA 500, 1000 and 1500 bp were used. The helicase assays were performed using the concentrations above but without surface immobilization or magnetic beads. The end-point fluorescence signal was then recorded after 5 min. This time should enable complete unwinding of the DNA substrate. The fluorescence signal can then be plotted against DNA base pairs to provide a signal calibration. Figures were made using PlotsOfData (Postma and Goedhart, 2019).

#### MDCC-SSB interactions with surface-immobilized dsDNA

1 mg Protein G dynabeads (2.8 μm) were prepared and loaded with 10 μg Anti-Digoxigenin antibody (Abcam ab6212) according to the manufacturer’s instructions. The Dynabead-Antibody was incubated with 1 μg 1500 bp DNA substrate for 15 min at room temperature. The streptavidin-coated surface was treated with an equivalent of 2 nM biotinylated 1500 bp DNA containing a 3’ digoxigenin for 15 min at room temperature in 50 mM Tris-HCl(pH7.5) and 150 mM NaCl. The surface was then washed three times with buffer. 100 nM MDCC-SSB was then added to the DNA and allowed to incubate for 10 min at room temperature. The MDCC fluorescence (ex. 430nm and em. 470 nm) was then recorded in the presence of the magnetic lid.

#### Live-cell experiments with the UAS-GAL4 reporter system

The UAS-Gal4 reporter U2OS cells, which stably expressed SNAP-hN1-Gal4 and the H2B-mCherry fluorescence reporter sequence, were cultured in McCoy’s 5A medium, supplemented with 10% FBS, 50 units/mL of penicillin and 50 µg/mL of streptomycin, and were maintained in a humidified incubator, at 37°C and in 5% CO2. Expression of the Notch receptor was induced by incubation with 2µg/ml doxycycline in cell culture medium for 24h, before use. Prior to measurements, the cells were first labelled with 5 µM biotin-conjugated SNAP-tag substrate (NEB) in tissue culture medium, by incubation for 30 min, at 37°C and 5% CO2. Cells were washed three times with medium and further incubated for 30 min in fresh medium at 37°C and 5% CO2. Medium was further replaced and supplemented with 30 µg MyOne Streptavidin C1 Dynabeads (1 μm) per condition. Cells were incubated for 30 min and then recording of mCherry fluorescence was performed in plate reader incubated at 37°C with 5 % CO2, over a period of 14h, in the presence of the magnetic lid. Due to potential cell-to-cell variation, the scan area was expanded to a 2 mm diameter circle consisting of 30 measurements.

### Graphics

Unless stated, data fitting and plotting was performed using Plots of data (Postma and Goedhart, 2019) and Grafit Version 5 (Erithacus Software Ltd).

## ACKNOWLEDGEMENTS

We thank Cancer Research UK (A26206), the MRC (MR/M020606/1) and Royal Society (RG150801) for funding.

## AUTHOR CONTRIBUTIONS

ADS, NF and CPT conceived the experiments. ADS, NF, DSP, YHG and CPT performed experiments and analysed data. ADS, NF and CPT wrote the manuscript.

